# Social behaviour and vocalizations of the tent-roosting Honduran White Bat

**DOI:** 10.1101/2021.03.01.433334

**Authors:** Ahana Aurora Fernandez, Christian Schmidt, Stefanie Schmidt, Bernal Rodríguez-Herrera, Mirjam Knörnschild

## Abstract

Bats are highly gregarious animals, displaying a large spectrum of social systems with different organizational structures. One important factor shaping sociality is group stability. To maintain group cohesion and stability, bats often rely on social vocal communication. The Honduran white bat, *Ectophylla alba* exhibits an unusual social structure compared to other tent-roosting species. This small white-furred bat lives in perennial stable mixed-sex groups. Tent construction requires several individuals and, as the only tent roosting species so far, involves both sexes. The bats’ social system and ecology render this species an interesting candidate to study social behaviour and social vocal communication. In our study, we investigated the social behaviour and vocalizations of *E. alba* in the tent by observing two stable groups, including pups, in the wild. We documented 16 different behaviours, among others, play and fur chewing, a behaviour presumably used for scent-marking. Moreover, we found 10 distinct social call types in addition to echolocation calls, and, for seven call types, we were able to identify the corresponding behavioural context. Most of the social call types were affiliative, including two types of contact calls, maternal directives, pup isolation calls and a call type related to the fur-chewing behaviour. In sum, this study entails an ethogram and describes the first vocal repertoire of a tent-roosting phyllostomid bat, providing the basis for further in-depth studies about the sociality and vocal communication in *E. alba*.

## Introduction

Bats are social animals exhibiting a large spectrum of social systems with varying degree of complexity. This includes species living in perennial stable groups)[e.g. 1] and species exhibiting social structures characterized by fission-fusion dynamics [e.g. 2]. One of the factors shaping sociality in bats is social group stability. Stable group living offers various benefits, including information transfer about food and roosts, and the evolution of cooperative behaviours such as allo-maternal care, allo-grooming and food sharing [2]. In bats, social vocal communication is a major factor facilitating group formation and cohesion thus supporting group stability [1, 3, 4]. A well-studied example, evolved to maintain group cohesion, is the contact call system in the foliage roosting bat, *Thyroptera tricolor* [5]. This bat species roosts in furled leaves of *Heliconia* plants, an ephemeral and often sparsely available resource. Roosts are only inhabitable for one day, hence, *T. tricolor* is daily forced to find and switch to new roosts [6]. Interestingly, despite constant roost switching, *T. tricolor* forms very stable perennial social groups [5]. To maintain group cohesion *T. tricolor* evolved a specialized call-and-response system, including inquiry calls to locate group members and response calls to recruit group members to the roost [6]. Another foliage roosting species, which exhibits an interesting social system is the phyllostomid bat *Ectophylla alba* [7]. *Ectophylla alba* is a small (i.e. 6-9g) neotropical phyllostomid bat species which is endemic to the Caribbean slope of Central America, known to construct and roost in tents [7, 8]. It is well known for its’ characteristic white fur and yellow skin colouration of the ears and nose-leaf, an adaption primarily evolved for camouflage associated with tent-roosting [9]. However, the yellow skin colouration, particularly of the nose-leaf, appears to be a sexually dichromatic trait, suggesting a secondary function as sexually selected signal [10]. The most commonly described mating system in tent-roosting bats is polygyny [i.e. harem structure composed of one male and several females; 11]. In contrast, *E. alba* forms mixed-sex groups with an average size of 5-6 individuals [12]. Interestingly, although genetic relatedness among adult social group members is very low [13], the groups are very stable over time, switch roosts together, and group members appear to have preferred individuals with whom they associate while roosting [12]. Furthermore, both sexes are involved in tent construction [8], in contrast, it is commonly supposed that in other tent-roosting speciesonly males construct tents [7]. Several individuals are involved in tent construction, a process which requires several nights until finalization [14]. However, it is not yet understood if these individuals all belong to the same social group.

*Ectophylla alba’s* unusual social organization, the high group stability despite the low relatedness and potential cooperative tent construction render this bat species an interesting candidate to study social behaviours and vocalizations mediating social interactions. Although the ecology of *E. alba* was intensively studied during the past decades, only very little is known about its’ social behaviour [12] and information about vocal communication is restricted to a single study, describing one call type emitted on the wing close to the roost [15]. This study aimed to describe the social behaviours and vocalizations in the roost to establish an ethogram and a vocal repertoire of *E. alba* based on observations of wild individuals in their natural habitat.

## Material and methods

### Study site and subjects

We monitored two groups of *E. alba* in La Tirimbina Biological Reserve, Heredia Province, in the North-East of Costa Rica (10°26’N, 83°59’W) from May to July 2010. La Tirimbina Biological Reserve contains fragments of secondary tropical wet forest and has been the centre of detailed investigations on the natural history of *E. alba* in the last decade [8, 10, 12, 14, 16-18]. The first *E. alba* group that we monitored consisted of four individuals (two adult males, one adult lactating female and her non-volant male pup), the second group (Fig 1 A) of 6-10 adult individuals of both sexes (the core group consisted of five adult males, one adult lactating female and her non-volant male pup; the sex of the other three adult bats that joined the core group on and off could not be determined). Females are polyestrous, they give birth to a single pup in September, respectively April [11]. Pups are born with fur and become more independent at an age of 3-4 weeks when they start to fly [14]. Bats of the first group could be individually distinguished via colour marks on their fur (Fig 1 B). Therefore, this was our focal group for behavioural observations and sound recordings; the other group was only occasionally observed and recorded to complement our data. During the 8-week observation period, both groups constructed new tents in the vicinity of their old ones and subsequently switched roosts.

**Fig 1.**
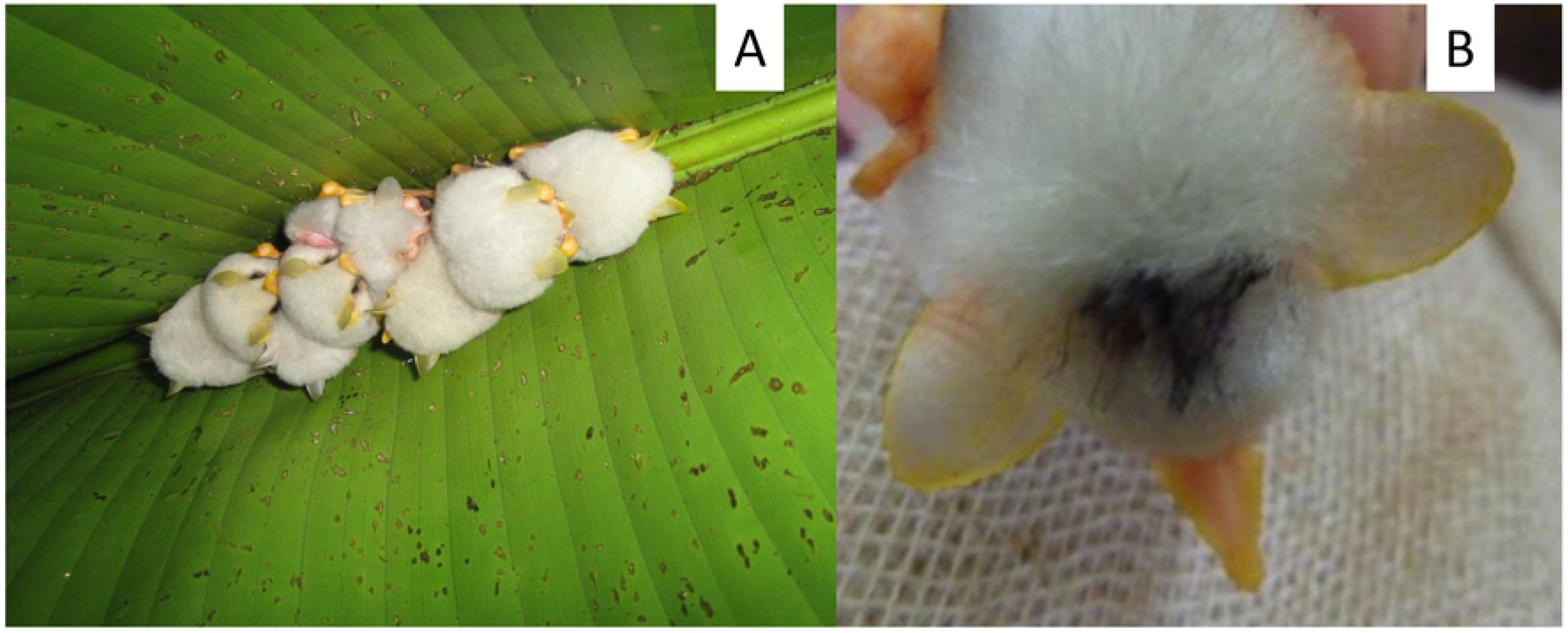
Observation of wild *Ectophylla alba*. **A:** Picture of the second group roosting in the tent during the day. Clearly visible is the yellow coloration of the ears of the adult individuals, whereas the pups’ ears are still almost white. **B:** Temporal colour marking of the fur to distinguish individual bats.

### Behavioural Observations

We conducted behavioural observations in the bats’ tents using a digital video camera with night-shot function (Sony Handycam DCR-SR32) and two infra-red lights (Sony HVL-IRM). The video equipment was placed directly underneath the tent and did not seem to disturb the bats. The video camera was connected via a 5 m cable to a video walkman (Sony DV-D900E) which allowed us to store the video recordings on mini-DV tapes (Sony DVM60PR3; 1.5 h run-time) and watch the video footage in real-time without disturbing the bats. This set-up permitted synchronous behavioural observations and sound recordings (see below for details). Video footage was analysed using the VLC Media Player (v1.0.5, VideoLan, France).

Since *E. alba* roosts cryptically during the day [7], we assumed that most social interactions would take place at dawn and dusk when bats are returning to or leaving the roost, or during the night when bats return to their tent [12]. To test this, we collected behavioural data over 22 hours by monitoring the first group for 3h periods spread equally over several days and nights covering the period from 9:47 am until 8:03 am of the next day. After defining the periods with the highest activity, we restricted our behavioural observations and sound recordings to these periods using *ad libitum* sampling [19]. Data was collected every night if the weather permitted it. During heavy rain, no data was collected.

Each social behaviour type was considered either a state or an event [19] and was listed in an ethogram. States were defined as behaviours with a minimum duration of ten seconds, including behaviours during which the same motor actions were performed repeatedly (e.g., wing fluttering). Events were instantaneous and singular (e.g. wing stretching) and occurred during a state. For each of the adult individuals in the first group (n=3) the 22 hours of observation time was split into two biological meaningful periods: the night-period, which included the time from leaving the tent at dusk to returning at dawn, and the day-period, which included the time from returning to the tent at dawn to leaving at dusk. Subsequently, we calculated the duration of each state (in seconds) and converted the durations into percentages to be able to compare these between individuals and day-respectively night-periods. The pup was not yet weaned and not foraging on his own; therefore, we decided not to split the 22-hour observation period for the pup.

### Sound Recordings

We used a high-quality ultrasonic recording setup (500 kHz sampling rate and 16-bit depth resolution) consisting of an ultrasonic microphone (Avisoft USG 116Hm with condenser microphone CM16; frequency range, 1-200 kHz) connected to a laptop computer (JVC, MP-XP741DE) running the software Avisoft-Recorder v4.2 (R. Specht, Avisoft Bioacoustics, Berlin, Germany). The behavioural context of each vocalization type in the vocal repertoire was assessed with simultaneous behavioural observations and video recordings. Sound and video recordings were synchronized with a bat detector (Pettersson D980, Pettersson Elektronik, Uppsala, Sweden) that was set on frequency division mode and connected to the audio channel of the video camera.

On one occasion, the video recordings contained two previously unknown vocalization types which were only recorded with the camcorder’s built-in microphone and not with the high-quality ultrasonic recording setup. Therefore, these two vocalization types were excluded from our acoustic analyses but we discuss the behavioural context in the results and corresponding spectrograms can be found in the supplements.

### Acoustical Analyses

Prior to acoustic analyses, vocalizations were visually classified into distinct social call types (i.e., social refers to calls other than echolocation calls) based on spectro-temporal features in the spectrograms; the different behavioural contexts in which vocalizations occurred were annotated based on behavioural observations of bats in the roost. Subsequent acoustic analyses were conducted to characterize the different social call types and assess their acoustic distinctiveness. We used Avisoft-SASLab Pro (v5.0, R. Specht, Berlin, Germany) for acoustic analyses. Only calls with good signal-to-noise ratio that were not overlapped by other calls or background noise were selected for acoustic measurements (116 in total; 5-41 per vocalization type). All calls were multiharmonic and some had an undulating structure (more than one frequency modulation). Since different harmonics were emphasized (i.e., had the largest amplitude) in different social call types, we used the strongest harmonic for measurements; we thus selected harmonics that contributed most to the acoustic impression of different social call types. We determined the start and end of calls manually based on the oscillogram. Subsequent measurements were taken from spectrograms created with a Hamming window with 512-point fast Fourier transform and 93.75 per cent overlap (frequency resolution: 977 Hz; temporal resolution: 0.064 ms). For all calls, we measured one temporal parameter (duration) and three spectral parameters (peak frequency at the start, middle and end of each call). Thus, we used four parameters per call to assess the acoustic distinctiveness of different vocalization types. Additionally, we further characterized undulating calls by measuring the peak frequency (pf) at every local maximum, minimum and inflexion point; values were subsequently averaged per call (mean maximum pf, mean minimum pf, mean inflexion point pf).

### Statistical analysis

We simultaneously included all four acoustic parameters in a discriminant function analysis (DFA), all of which were checked for multicollinearity. We used an ‘n-1’cross-validation procedure which classified each call based on discriminant functions established with all social call types other than the one being classified. Prior probabilities were adjusted to unequal group sizes. All statistical tests were performed with SPSS (v.22, SPSS Inc., Chicago, IL, U.S.A.)

## Results

### Social behaviours

We observed six behavioural states and ten different behavioural events occurring in the tent-roost (Table 1). Most behaviours were observed during both the day-and night-period. During the 22-hour observation period, the states “fur chewing” and “playing” were only observed during the night-period, whereas the event “change position” was only noted during the day-period. However, outside of the 22-hour observation period, “change position” was also observed during the night.

**Table 1.** Ethogram describing behaviours of *E. alba* observed in the tent.

| State or Event | Behaviour | Description | Estimated frequency of occurrence | Performing sex | Age |
| --- | --- | --- | --- | --- | --- |
| S | Resting | Roosting motionless, often with concealed faces (not to be equated with sleeping) | Very common | Both sexes | all |
| S | Scanning | Echolocating with twitching ears and nose-leaf, often directed at the ground below the roost | Very common | Both sexes | All |
| S | Self-Grooming | Tending to coat and wings with tongue and hind feet | Very common | Both sexes | All |
| S | Fur chewing | Conspicuous, prolonged licking and/or gentle biting the fur of conspecifics (of both males and females and pups) between the shoulder blades and coating it with saliva | Rare | Only males | Adults |
| S | Allo-Grooming | Mother grooming her non-volant pup | Common | Only females | Adults |
| S | Nursing | Mother breastfeeding her non-volant pup | Common | Only females | Adults |
| S | Licking | Soliciting maternal care by licking the corners of her mouth or her belly; often followed by nursing | Common | Both sexes | Pups |
| S | Twitching | Maternal signal for the pup to release the teat | Common | Only females | Adults |
| S | Shaking | Rapid whole-body muscle contractions, presumably for thermoregulation | Rare | Both sexes | Pups |
| S | Playing | Conspicuous, prolonged and seemingly playful engagement with a torn piece of leaf in the roost | Rare | Both sexes | pups |
| S | Startle posture | Raising of half-opened wings high above head and back, followed by covering face with raised half-open wings | Rare | Both sexes | Pups |
| S | Flight practice | Rapid wing fluttering while clinging to the roost surface with hind feet | Common | Both sexes | Pups |
| E | Yawning | Exposing gum and teeth (when resting, cleaning or nursing) | Common | Both sexes | All |
| E | Changing position | Climbing to a different roosting position within the day-roost (when resting) | Very common | Both sexes | All |
| E | Wing stretching | Stretching of one wing (when resting, cleaning or nursing) | Very common | Both sexes | All |
| E | Hitting | Aggressively hitting conspecifics with partly outstretched wing (when resting) | Rare | Both sexes | Adults |
| E | Urinating/Defecating | Urinating/Defecating by arching back; never when in body contact with conspecifics (when cleaning) | Rare | Both sexes | All |

Adult bats exhibited two main activity peaks during the 22 hours; one before sunset and one around sunrise. This coincides with the time at which adult bats leave, respectively, return from foraging at night. The activity peaks were characterized by increased auto-grooming, wing stretching and frequent position changing in the roost. The analysis of the 22-hour observing period revealed that during the day-period, the adult bats spent the majority of time resting (male 1: 91.9%, male 2: 96.7%, female: 77.4% of the time). A short amount of the time they spent auto-grooming (male 1: 8.1%, male 2: 1.5%, female: 1.5% of the time) and, in the case of the female, nursing (21.1%). The female nursed the pup three times before leaving for foraging at sunset and twice in the early morning after returning to the tent at sunrise. The longest nursing duration was observed at 6 am when the female nursed her pup for one hour and 21 minutes.

During the night-period, adult bats spent most of the time foraging (i.e., were absent; male 1: 96.5%, male 2: 95.2 %, female: 95.9% of the time). The female visited the tent twice during the night to nurse her pup (3.6%) and spent some time self-grooming (0.5%). One of the males visited the tent only once, whereas the second male visited the tent five times and stayed for a short period. During their visits, males were resting (male 1: 0.3%; male 2: 0.8%) and self-grooming (male 1: 1.08%, male 2: 2.5%). Furthermore, they engaged in fur chewing (male 1: 2.14%, male 2: 1.4%).

The pups’ main activities (22h-period was not split in day- and night-period) constituted of resting (63.7%), nursing (12.8%), auto-grooming (8.5%) and changing position in the tent (0.9%). Although changing position was usually considered an event, in this case, it was defined as a state because the pup was continuously changing his position in response to being gently bitten by an adult male (i.e., male fur chewing behaviour). At night, the pup was absent for short periods of time (14.1%). At this stage in ontogeny, the pup engaged in first flight attempts. However, compared to the other group members, the pup was the least time away from the tent. Furthermore, the pup occasionally engaged in a behaviour which was defined as play (Table 1). Note that some states were not observed during this continuous 22-hour recording, therefore, they are not included in the calculation of time-budgets but are nevertheless described in the ethogram. Two unusual and rare behaviours are subsequently described in greater detail; namely “fur chewing” and “play”.

#### “Fur chewing” in males

After manoeuvring behind the back of a roosting group member, males were observed licking and gently biting the fur between the shoulder blades for a prolonged time (up to 13 minutes, Fig 2). While chewing the fur, males sometimes simultaneously were trembling their folded wings. After chewing, the bitten individual showed a visible patch of wet fur from the saliva. In most cases, males were chewing fur on the back of a female, but it was also observed that males chewed on the back of each other. In one occasion, a male that returned to the roost performed this behaviour on the pup who was roosting alone in the tent (for about 6 minutes, see video S1). The individual being bitten remained mostly calm, sometimes started self-grooming, wing stretching and changing the position, with the fur chewing individual firmly clinging on. Eventually, the individual being bitten (if it was not the pup) also engaged in biting/licking another group member.

#### “Play” in pups

While being alone in the tent, the pup started to investigate a torn piece of the roost leaf (see video S2, Fig 2). First, the pup started sniffing the leaf piece, and soon after used both thumbs/claws and wrists to grasp the leaf piece. Once grasped, the pup started chewing on the piece. The pup chewed on the leaf piece for a few seconds, stopped and started scanning. This behavioural sequence was repeated several times. Sometimes, the pup also started cleaning, wing stretching or moving around after chewing on the leaf piece. At one time, the pup was observed to inspect the modified midrib of the tent (i.e. the part of the leaf which is modified during tent construction to collapse the leaf next to the cut to achieve the typical shape of *E. alba* tents, see [8]. Afterwards, the pup turned back to the leaf piece and started chewing again, while simultaneously using the wrists and claws grasping and holding on to it. Chewing could get quite vigorous, and eventually, the pup started to bend the leaf piece to some extent.

**Fig 2.**
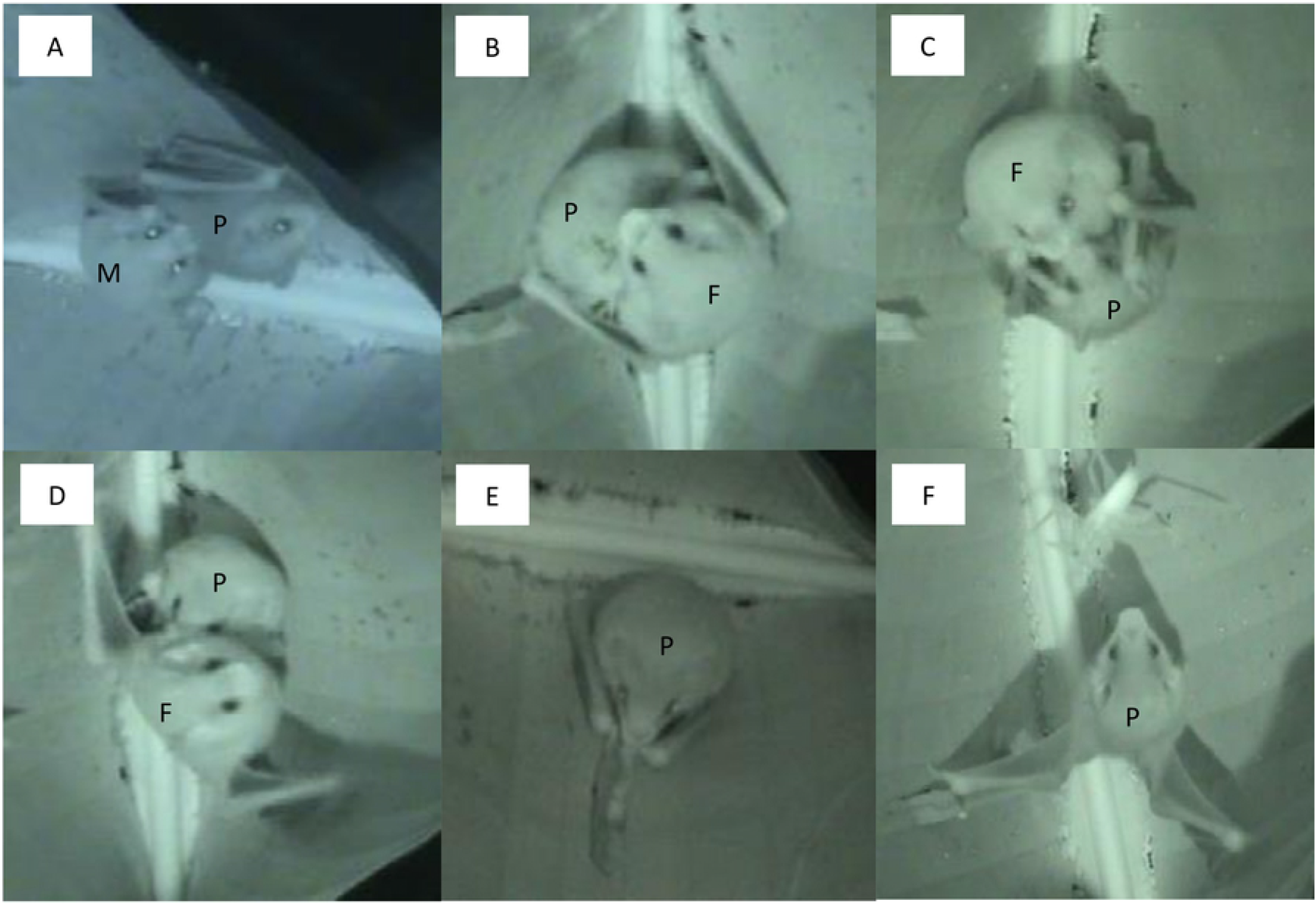
Social behaviours of *E. alba*. **A:** Fur chewing behaviour. The male (M) is chewing/licking the back of the pup (P). **B:** Nursing. The pup (P) is attached to the nipple of the female (F). **C:** Licking. The pup (P) is licking the mouth of the female (F) to solicit nursing. **D:** Twitching. The female (F) is shaking the pup (P) off after nursing. **E:** Play. The pup is playing with a torn leaf piece inside the tent. **F:** Startle response. The pup (P) shows the typical wing posture when frightened, in this case by a grasshopper that has wandered into the tent. For detailed descriptions of the behaviours see Table 1.

### Social vocalizations

*E. alba* produced ten distinct social call types in addition to echolocation calls, and for most call types the behavioural context in which they were uttered could be defined. The social call types SC9 and SC10 were not included in the statistical analysis because they were only recorded once with the microphone of the camcorder (Fig S1). The visual classification of eight social call types was confirmed by the classification success of the cross-validated DFA (88.8% of all call types were classified correctly, Table 2, Fig 3B). The acoustic parameter that contributed most to the distinction of social call types was peak frequency in the centre of the call, peak frequency at the start and the end of the call contributed moderately to the call distinction, whereas duration only played a minor role. A conspicuous feature of the social vocalizations in *E. alba* is the suppression of the fundamental frequency and the lower harmonics in some of the call types (SC1, SC2; SC5-SC7, Table 3).

**Table 2.**
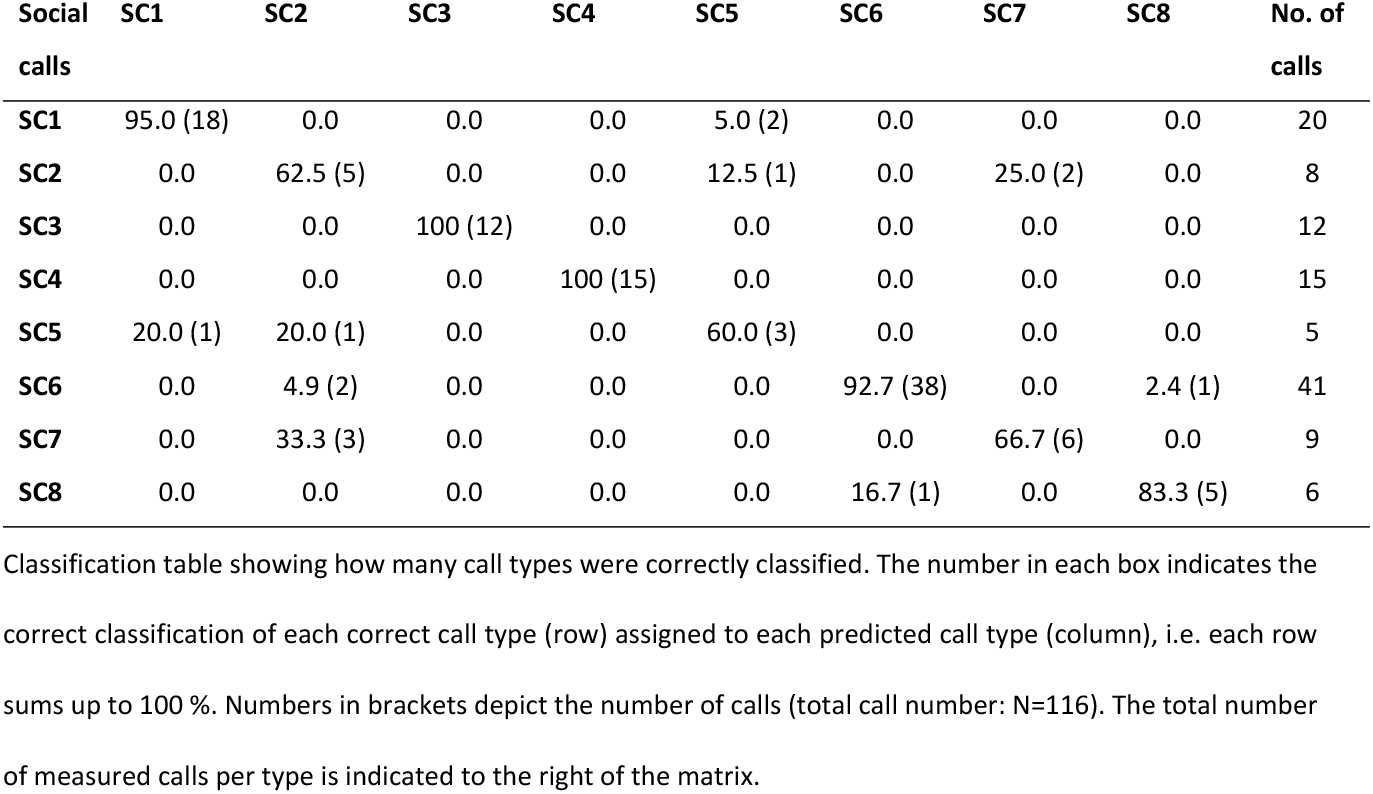
Classification success in per cent (%) of the cross-validated discriminant function analysis.

| Social calls | SC1 | SC2 | SC3 | SC4 | SC5 | SC6 | SC7 | SC8 | No. of calls |
| --- | --- | --- | --- | --- | --- | --- | --- | --- | --- |
| SC1 | 95.0 (18) | 0.0 | 0.0 | 0.0 | 5.0 (2) | 0.0 | 0.0 | 0.0 | 20 |
| SC2 | 0.0 | 62.5 (5) | 0.0 | 0.0 | 12.5 (1) | 0.0 | 25.0 (2) | 0.0 | 8 |
| SC3 | 0.0 | 0.0 | 100 (12) | 0.0 | 0.0 | 0.0 | 0.0 | 0.0 | 12 |
| SC4 | 0.0 | 0.0 | 0.0 | 100 (15) | 0.0 | 0.0 | 0.0 | 0.0 | 15 |
| SC5 | 20.0 (1) | 20.0 (1) | 0.0 | 0.0 | 60.0 (3) | 0.0 | 0.0 | 0.0 | 5 |
| SC6 | 0.0 | 4.9 (2) | 0.0 | 0.0 | 0.0 | 92.7 (38) | 0.0 | 2.4 (1) | 41 |
| SC7 | 0.0 | 33.3 (3) | 0.0 | 0.0 | 0.0 | 0.0 | 66.7 (6) | 0.0 | 9 |
| SC8 | 0.0 | 0.0 | 0.0 | 0.0 | 0.0 | 16.7 (1) | 0.0 | 83.3 (5) | 6 |

Classification table showing how many call types were correctly classified. The number in each box indicates the correct classification of each correct call type (row) assigned to each predicted call type (column), i.e. each row sums up to 100 %. Numbers in brackets depict the number of calls (total call number: N=116). The total number of measured calls per type is indicated to the right of the matrix.

**Table 3.** Acoustic parameters of the eight social call types.

| Social calls | N | Duration [ms] | Peak freq. start [kHz] | Peak freq. centre [kHz] | Peak freq. end [kHz] | Measured harmonic |
| --- | --- | --- | --- | --- | --- | --- |
| SC1 | 20 | 5.01 ± 1.2 | 116.6 ± 13.0 | 79.8 ± 7.8 | 57.6 ± 6.2 | 3d |
| SC2 | 8 | 9.6 ± 4.8 | 94.3 ± 4.7 | 83.5 ± 4.2 | 80.2 ± 5.8 | 10th |
| SC3 | 12 | 50.5 ± 6.5 | 21.0 ± 0.6 | 20.4 ± 1.3 | 20.5 ± 1.6 | 1st |
| SC4 | 15 | 24.4 ± 6.5 | 41.3 ± 2.6 | 39.7 ± 1.5 | 40.0 ± 2.3 | 2nd |
| SC5 | 5 | 7.1 ± 2.4 | 99.5 ± 19.2 | 72.3 ± 5.9 | 65.9 ± 3.9 | 3th |
| SC6 | 41 | 49.1 ± 20.4 | 86.3 ± 8.8 | 84.3 ± 6.3 | 86.3 ± 8.3 | 8th |
| SC7 | 9 | 9.3 ± 2.8 | 93.6 ± 11.7 | 91.6 ± 10.2 | 93.1 ± 13.8 | 4th |
| SC8 | 6 | 18.9 ± 4.5 | 83.0 ± 9.1 | 87.7 ± 11.2 | 60.6 ± 11.9 | 3th |

The table depicts mean and standard deviation for each social call type averaged over all calls measured per type (column 2). The frequency parameters were measured in the harmonic which contained the most energy (column 7).

**Fig 3.**
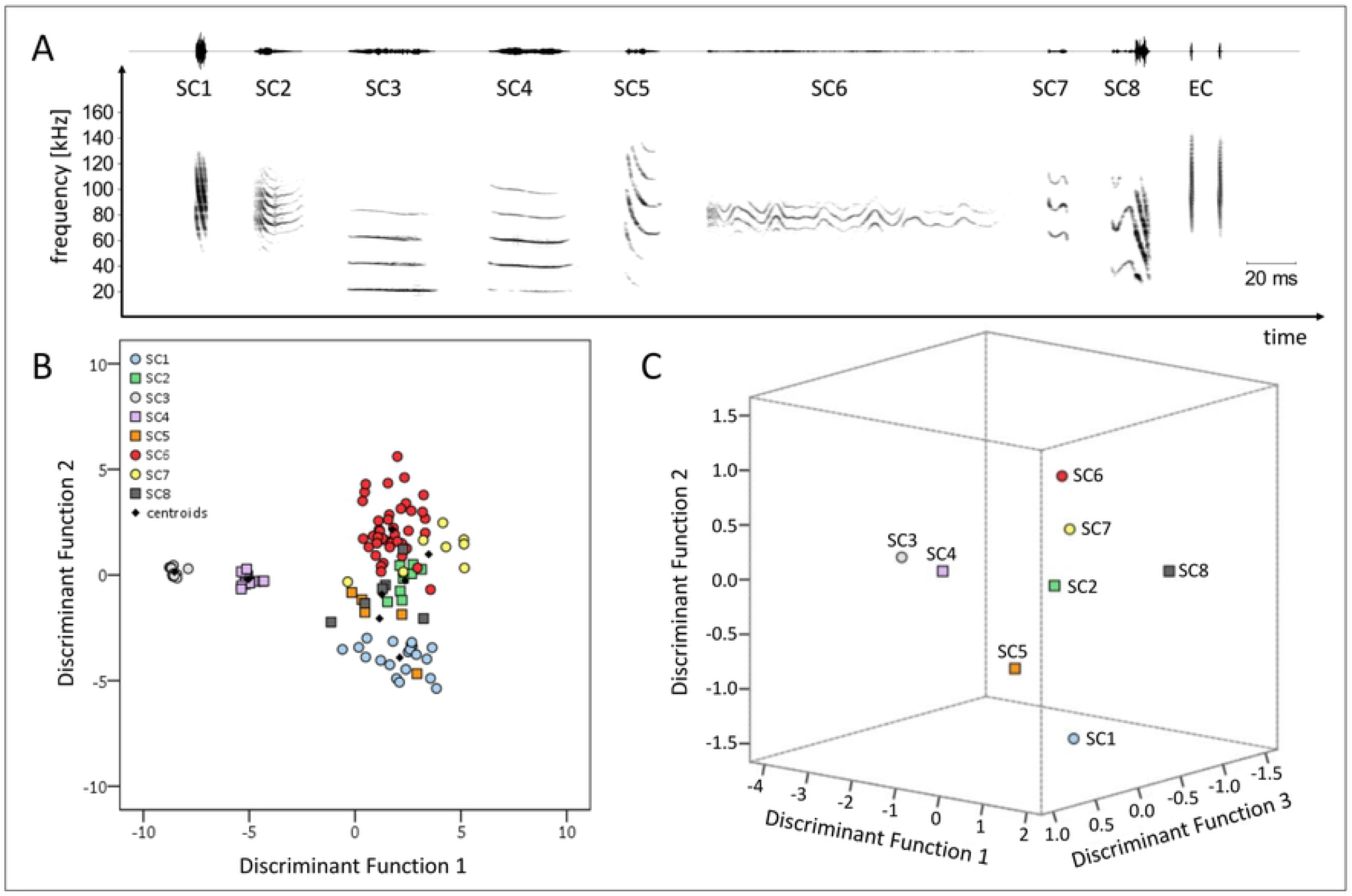
The eight social call types of *E. alba*. **A:** Spectrograms depicting the eight social call types (SC1-SC8) and two echolocation calls (EC) of *E. alba*. The spectrograms correspond to the natural appearance of those call types; i.e., suppression of the lower harmonics in types SC2, SC5, SC6, SC7. Social call types SC9 and SC10 are depicted in the supporting information (S_Fig 1). Information about acoustic parameter measurements is depicted in Table S1. Echolocation calls are shown for comparative reasons. The context in which the social call types were emitted is described in Table 2. Spectrograms were created using a 1024-point FFT and a Hamming window with 87.5 % overlap. **B:** The spacing of the eight social call types of *E. alba* in a two-dimensional signal space defined by the first 2 discriminant functions. Each social call type is represented by a distinct symbol, small black symbols represent centroids (i.e., the canonical mean of all calls per type). Note that EC are not included in the DFA.

The DFA included acoustic measurements obtained from the harmonics that contributed most to the acoustic impression of different social call types. As a control, we calculated a second DFA including acoustic measurements of the fundamental frequency which corroborated the results of the first DFA (supporting information).

Furthermore, for seven out of ten social call types the behavioural context in which they were uttered was elucidated (Table 4). Three social call types were uttered in an affiliative context, namely SC2, SC3, SC5 (Fig 3). Two social call types were uttered in the context of mother-pup interactions, namely SC4 and SC6 (Fig 3). In agonistic context, two social call types were uttered (see supplements), namely SC9 and SC10. Echolocation calls were uttered during flight and during alert behaviour in the roost (i.e., scanning, see Table 1). Most social call types were uttered singly and with a single exception (SC7) monosyllabic (Table 4).

**Table 4.** Behavioural context of the social call types.

| Social call | Production mode | Syllable type | Behavioural context | Frequency of occurrence (ranked) | Sex | Age |
| --- | --- | --- | --- | --- | --- | --- |
| SC1 | In series | Monosyllabic | Unknown | 1 | Both sexes | All |
| SC2: Grooming/biting call | Singly | Monosyllabic | Emitted while male gently bites or grooms the neck of a female | 2 | Males Females? | Adults |
| SC3: Contact call 1 | Singly | Monosyllabic | Emitted on the wing by individuals flying in the vicinity of the tent (both empty and occupied tent), and emitted while leaving the tent | 3 | Unknown | Adults |
| SC4: Maternal directive | Singly | Monosyllabic | Emission immediately followed by nursing | 2 | Females | Adults |
| SC5: Contact call 2 | Singly | Monosyllabic | Emitted on the wing by individual flying in the vicinity of the tent; and emitted while leaving the tent | 4 | Unknown | Adults |
| SC6: Isolation call | Singly | Monosyllabic | Solicitation of nursing | 2 | Both sexes | Pups |
| SC7 | Singly | Multisyllabic | Unknown | 5 | Unknown | Unknown |
| SC8 | Singly | Monosyllabic | Unknown | 2 | Unknown | Unknown |
| SC9 | In series | Monosyllabic | Aggressive/Distress | 6 | Both sexes | Unknown |
| SC10: Screech | In series | Monosyllabic | Aggressive/Distress | 6 | Both sexes | Adults |

The frequency of occurrence corresponds to the number of recording sessions during which the social call type was recorded: 1 being the most frequent type, 6 the less frequent type. Sex denotes the sex of the emitter of a given social call type. Age denotes which age group (pup/adult) uttered the social call type.

## Discussion

This study provides the first description of the behavioural ethogram and vocal repertoire of *E. alba* in the roost. We identified 16 different behaviours, including two particularly interesting ones; the “fur chewing” behaviour performed by adult males, and, the description of a pup behaviour, which very likely meets the criteria defining play in animals. The vocal repertoire is constituted of 10 distinct social call types, and for seven call types, the behavioural context was determined.

The temporal occurrence of social behaviours shows clear differences between day and night periods. During the day, aside from the two main activity peaks at dusk and dawn, the bats were almost exclusively resting. Roosting quietly during the day could be owned to avoid alerting day predators, such as primates [20, 21] but could also be a method to save energy. At night, as expected, adult individuals spent most of their time foraging but paid short visits to their tent. During her visits at night, the female was regularly nursing her pup. Former research also showed that, during their visits at night, females spent a considerable amount of time nursing and/or grooming the pup, especially during early ontogeny before pups became more independent [12]. Because pups are born almost furless, maternal care is especially crucial during the first days after birth, probably for reasons of thermoregulation. Adult males mostly engaged in self-grooming during their roost visits at night. Furthermore, the most interesting behaviours, “fur chewing” performed by adult males and “play” in pups, were both only observed at night. As described in the results, “fur chewing” was performed by males only. Our observations conform to the description of the behaviour in a former study, by Rodríguez-Herrera and colleagues [12]. A new observation in our study was that adult males perform “fur chewing” not only on adult males and females but also on pups. From the behavioural response of the bitten individuals, both adult and pup, it seemed that “fur chewing” did not cause any pain (see also Rodríguez-Herrera et al. 2019) [12]. At most it was possibly perceived as irritating (e.g., for a while, the pup who was being bitten tried to shake off/get away from the male, see video S1) as the bitten individual hardly showed any serious attempts of escape or strong resistance.

It remains speculative if “fur chewing” is a behaviour for scent-marking group members or if it is a form of allo-grooming. Allo-grooming is mainly observed in stable social groups (e.g. harems, maternity colonies) with varying degree of relatedness [2]. Besides strengthening of social relationships, allo-grooming is also exchanged for other social benefits [1, 2, 22, 23]. So far, allo-grooming among adult individuals (not including mother-pup grooming) was described for a few bat species only [2, 24]. However, in contrast to other species where allo-grooming was observed [e.g. 1, 22, 23], *E. alba* seems to restrict grooming to a very specific spot on the back, not including other body parts [this study and 12]. Furthermore, after “fur chewing”, a visible wet spot remained on the back of the receiver (video S1). This is reminiscent of the scent-marking behaviour of group members in *Noctilio leporinus* and *Cynopterus sphinx* [25, 26]. *Noctilio leporinus* females, who form stable perennial groups, rub their heads on other females’ heads and backs to scent-mark them [25]. In *C. spinx*, individuals form so-called grooming clusters, where individuals hold on to each other while distributing saliva on body parts of group members [26]. In both species, scent-marking was associated with group member recognition. In *E. alba*, “fur chewing” could have a similar function; group recognition through scent. *Ectophylla alba* forms very stable social groups [12], although the genetic relationship between adult individuals is almost zero [13]. It is known that groups switch together to new roosts, which are usually close to the currently occupied tent within a small area [17]. Tent construction is costly; time spent manipulating a leaf cannot be invested in foraging, and construction requires several nights [14]. Furthermore, *E. alba* has specific requirements to its roosting microhabitat [20], probably limiting the availability of potential roosting areas and, therefore, increasing the value of suitable places. A scent signature could assist the identification and recognition of social group members joining a roost. This would also explain why males perform this behaviour not only on females but also on pups. Nevertheless, besides scent-marking “fur chewing” could also strengthen social bonds between individuals, as observed in other species [2]. It remains to be investigated which function(s) the observed “fur chewing” plays in *E. alba*.

A behaviour characterized as play in animals is defined as (i) a non-fully functional behaviour, (ii) being spontaneous and voluntary, (iii) different from a formal performance of functional behaviour, e.g. exaggerated or incomplete, (iv) repeatedly performed during a period of an individual’s life, and, (v) performed only when the animal is free from stress [27]. There are several hypotheses about the function of play in young animals [e.g. 28], and despite opposing views regarding certain aspects, most agree that one of the main functions of play is to refine one’s motor skills [29]. Play behaviour is grouped into three categories; social play, locomotor play and object play [29]. Studies about play in animals are very scarce, and in bats, it has only been observed in a few occasions. Social play was described in *Pteropus giganteus*, where young individuals engage in play-fight and wrestling, first with their mothers, later among subadult individuals [30]. Similar play fighting was also described for other Pteropodidae species [31]. Young vampire bats engage both in object-and social play, the latter involving mounting, wrestling and chasing [32]. The pups’ handling of the torn leaf piece, we observed for the first time in this study, meets the criteria of object play behaviour. The pups’ object play might be a precursor to later actual tent construction behaviour. Adult tent construction involves biting and puncturing the leaf using the teeth, and further extension of these holes by claws until the leaf collapses next to the cut [8]. By grasping a part of the leaf with the thumbs and repeatedly pulling it up, down and inward the leaf bends downward forming the final shape of the tent [8]. Several motor patterns of adult tent construction are found in the pup’s object play; the chewing of the leaf, the grasping of a leaf piece with the thumbs and wrists and finally the bending and moving of the leaf piece (but instead of using the thumbs the pup used both, thumb and mouth, see video S2). The pups’ behaviour is reminiscent of motor patterns used by adults in tent construction, but not yet fully functional (criteria i & iii). The pup was free from stress, voluntarily engaging with the leaf piece (criteria ii & iv). Moreover, this behaviour was observed several times during different nights (v). However, our sample size is restricted to observations of a single pup. The observed behaviour could also be explained by curiosity towards an unexpected object (torn leaf piece) present in the roost.

The repertoire size of *E. alba* described here is within the size range of the vocal repertoire sizes of other phyllostomid bats (*Glossophaga soricina* [n=15 social call types, 33], *Glossophaga commissarisi* [n=8 social call types, 33], *Carollia perspicillata* [n=10 social call types, 34] and *Phyllostomus discolor* [n=12 social call types, 35]). Our description of the vocal repertoire is incomplete, because we did not observe courtship behaviour. Nevertheless, the repertoire described here most likely includes the majority of the social call types (outside of the mating context). Most of the social call types were affiliative (SC2-SC6), and although the exact context of call types SC1, SC7-SC8 was not elucidated, they were uttered in neutral, non-aggressive situations (Table 4). This corresponds to the observed behaviours; aggressive/agonistic behaviour like “wing hitting” was only rarely observed (Table 1). The only aggressive/distress vocalizations (Fig S1) were recorded during an incident when a mosquito stung a bat in a sensitive spot on his back that was temporarily hairless because of a telemetry tag that had recently fallen off. Similarly, a study from Rodríguez and colleagues never noticed agonistic behaviour in the group they observed [12]. So far, it is still unknown whether males exhibit aggressive behaviour in the mating context, and, for example jointly defend a tent or their roosting area.

The pup isolation call of *E. alba* is different from the isolation calls of other phyllostomid bats because its fundamental frequency and the lower harmonics are suppressed (Fig. 3, Table 3). In other phyllostomids studied so far, most of the sound energy of pup isolation calls is located in the fundamental frequency [P. hastatus, P. discolor, G. soricina, C. perspicillata 36, 37-39]. This particular spectral characteristic of *E. alba* isolation calls might be an adaption to its roosting ecology. The tents offer less protection from predation compared to the roosting sites of the other species, due to their resistance and stability (i.e. leaf versus cave or tree hole) and probably also due to their location (in the understory, less than 2m above ground) [14]. Restricting the isolation calls’ sound energy to a narrow high-frequency band could create a communication channel for *E. alba* that is circumventing the hearing range of some predators; for instance, it is known that small primates successfully predate tent roosting bats [21]. The isolation call of *E. alba* also differs in its duration from the isolation calls of the other species. Why these isolation calls have such a long duration and whether this is possibly related to the intensity of solicitations for nursing could be investigated in further studies.

The social call types SC3 and SC5 were uttered on the wing, in the vicinity to the roost, and while leaving the tent. The call type SC3 is similar to the social call described in the study of Gillam and colleagues [15]. They recorded this specific social call in the vicinity of tent roosts and once before entering the roost [15] which corroborates our observations. Following Gillam and colleagues, we hypothesise that these social call types serve as contact calls. They are not used to attract and recruit group members for roosting, a function that contact calls recorded in other foliage-roosting bats have [6, 15]. They might serve for the coordination of group formation; however, the purpose of group formation in the vicinity to the tent is completely speculative at this point. Besides the potential function of group formation, these calls might additionally signal roosting area occupancy to other social groups roosting in the vicinity. In both scenarios, group member recognition is crucial; therefore, in future studies, it would be interesting to investigate if the social call types SC3 and SC5 encode a vocal group signature and if they elicit phonotaxis in receivers.

The affiliative social call type SC2 was uttered before “fur chewing”, always by the active (i.e., fur chewing) and never by the passive (i.e., individual whose fur was chewed) bat. We never recorded a vocal response of the passive individual. To our knowledge, there is no other study describing a social call associated with scent-marking and/or allo-grooming. This social call might be an appeasement vocalization to signal non-aggressive intention towards the passive individual. Further, it might simultaneously strengthen dyadic relationships between individuals. *E. alba* seem to prefer to associate with particular group members within the roost [12], and this call might encode an individual signature facilitating social interactions. However, it is not known if “fur chewing” occurs more frequently between particular group members.

Overall, our research contributes to the growing number of studies on social behaviour and vocal repertoire descriptions in phyllostomid bats. Our results allowed us to raise new questions and formulate hypotheses about particular social behaviours and social call types which can be tested in future observational and experimental studies. With our study, we hope to initiate further studies about social behaviour not only in *E. alba* but other bat species, and especially encourage further studies describing vocal repertoires of bats to assess the communicative capacity of this speciose taxon.

## Acknowledgements

We would like to thank the entire team of the Tirimbina Biological Reserve for their support and providing excellent research conditions. Simon P. Ripperger helped to improve this manuscript with well-conceived comments.

## Supporting information

**S1 Video. Male fur chewing**. This video captures the pup and an adult male together in the roost at night (the other four individuals of this social group are absent). Directly after landing in the tent, the male briefly smells the pup and directs himself behind the pup. Immediately, he starts biting/chewing the pup’s fur. The biting is accompanied by wing trembling. The pup seems irritated, trying to move around. At some point, it attempts to stretch its wing. After a few seconds, the male starts cleaning himself. At the end of the video the pup turns around and a wet part can be spotted on its back.

**S2 Video. Pup play behaviour**.

This video captures the pup alone in the roost at night. The detailed description of the pup’s behaviour can be found in the results section of the study.

**S1 Fig. Spectrograms of social call types SC9 and SC10**.

Spectrograms depicting the social call types SC9 and SC10 of *E. alba* **(A)**. Both social call types were emitted by an adult male in response to a mosquito sting on a hairless spot on the back of the bat. The vocalizations were accompanied by agonistic behaviour, including wing flapping and hitting conspecifics with folded wings. Both social call types were emitted in series and both were concatenated to a vocal sequence **(C)**. Both social call types were recorded only with the camcorder’s built-in microphone and not with the high-quality ultrasonic recording setup. This is the reason why the recordings are clipped at 20 kHz. Spectrograms were created using a 512-point FFT and a Hamming window with 75% overlap.

